# Disruption of oncogenic targeting by ISWI via phosphorylation of a prion-like domain

**DOI:** 10.1101/2020.03.11.987750

**Authors:** Mark Chen, Joseph P. Foster, Ian C. Lock, Nathan H. Leisenring, Andrea R. Daniel, Warren Floyd, Eric S. Xu, Ian J. Davis, David G. Kirsch

**Affiliations:** Department of Pharmacology and Cancer Biology, Duke University Medical Center, Durham, North Carolina, 27708; Medical Scientist Training Program, Duke University Medical Center, Durham, North Carolina, 27708; Curriculum in Bioinformatics and Computational Biology, University of North Carolina at Chapel Hill, Chapel Hill, North Carolina, 27599; Department of Radiation Oncology, Duke University Medical Center, Durham, North Carolina, 27708; Lineberger Comprehensive Cancer Center, University of North Carolina at Chapel Hill, Chapel Hill, North Carolina, 27599; Departments of Pediatrics and Genetics, University of North Carolina at Chapel Hill, Chapel Hill, North Carolina, 27599

**Keywords:** Cancer, chromatin, ISWI, sarcoma, fusion oncoprotein, radiation biology, phase separation

## Abstract

Chromosomal translocations generate oncogenic fusion proteins in approximately one-third of sarcomas, but how these proteins promote tumorigenesis and the effect of cancer therapies on their function are not well understood. Here, we reveal a molecular mechanism by which the fusion oncoprotein FUS-CHOP promotes tumor maintenance that also explains the remarkable radiation sensitivity of myxoid liposarcomas. We identified novel interactions between FUS-CHOP and chromatin remodeling complexes that regulate sarcoma cell proliferation. One of these chromatin remodelers, SNF2H, co-localizes with FUS-CHOP genome-wide at active enhancers. Following ionizing radiation, DNA damage response kinases phosphorylate the prion-like domain of FUS-CHOP to impede these protein-protein interactions, which are required for transformation. Therefore, the DNA damage response after irradiation disrupts oncogenic targeting of chromatin remodelers required for FUS-CHOP-driven sarcomagenesis.

**Significance:** Prion-like domains translocated in cancer have been shown to drive global epigenetic changes that are oncogenic. However, some translocation-driven cancers exhibit dramatic clinical responses to therapy, though the mechanism for these responses are not well-understood. Here we show that ionizing radiation can disrupt oncogenic interactions between a fusion oncoprotein and a chromatin remodeling complex, ISWI. This mechanism of disruption links phosphorylation of the prion-like domain in an oncogenic fusion protein to DNA damage after ionizing radiation and reveals that a dependence on oncogenic chromatin remodeling underlies sensitivity to radiation therapy in myxoid liposarcoma.

## Introduction

Translocation-driven sarcomas, such as Ewing sarcoma, comprise around one third of soft tissue sarcomas (Mertens et al., 2016). In approximately half of these sarcomas, the N-terminal domain of the chimeric oncoprotein is derived from one of the FET proteins (FUS, EWSR1, TAF15). For example, in Ewing sarcoma, the EWSR1 N-terminus prion-like domain (PrLD) is fused to the carboxyl terminal domain of an ETS family transcription factor, typically FLI1, to generate EWS-FLI1. EWS-FLI1 acts as a neomorphic transcription factor by targeting GGAA-containing microsatellite repeats and altering chromatin accessibility mediated through interactions with the BAF (BRG1/BRM-associated factors) complex (Boulay et al., 2017; Lindén et al., 2019; Patel et al., 2012). Importantly, the interactions between EWS-FLI1 and the BAF complex are mediated through the PrLD of EWSR1(Boulay et al., 2017).

In a subset of Ewing sarcomas, FUS (fused in sarcoma) constitutes the N-terminal domain of the fusion protein(Shing et al., 2003). FUS is also the N-terminus partner of other fusion proteins that drive sarcomagenesis. FUS-CHOP (also known as FUS-DDIT3) results from a t(12;16)(q13;p11) translocation that drives 95% of myxoid liposarcomas (MLPS) (Aman et al., 1992; Antonescu et al., 2001; Crozat et al., 1993; Panagopoulos et al., 1994). Exome analysis of MLPS has revealed few other recurrent mutations in coding sequences(Joseph et al., 2014). FUS-CHOP is specific to MLPS and has not been detected in other liposarcoma subtypes or other cancers(Wylie, 2004). These data suggest that the FUS-CHOP translocation is the predominant driver in MLPS tumorigenesis, though its precise mechanism is unknown. Over a century ago, James Ewing recognized that, like Ewing sarcoma, these sarcomas with a myxoid background were exquisitely sensitive to radiation therapy(Ewing, 1935), but the mechanism underlying this radiosensitivity remains unknown. Indeed, MLPS is distinguished by its remarkable clinical response to radiation therapy compared to most other sarcoma subtypes (Chung et al., 2009; Guadagnolo et al., 2008; Pitson et al., 2004). We hypothesized that the PrLD of FUS-CHOP interacts with chromatin remodeling complexes and identified new interactions with imitation switch (ISWI) chromatin remodeling complexes. Furthermore, we hypothesized that disruption of these interactions contributes to the radiosensitivity of MLPS and determined that phosphorylation of the PrLD mediates these interactions, but impedes transformation by FUS-CHOP.

## Results

### FUS-CHOP interacts with the BAF and ISWI chromatin remodeling complexes

To identify chromatin remodeling complexes that interact with FUS-CHOP, we performed proximity ligation assays (PLA) for three of the major chromatin remodeler complex families: BAF, ISWI, and nucleosome remodeling deacetylase (NuRD). We found that all three chromatin remodeler families were in close proximity to FUS-CHOP in the nuclei of human MLPS cells (MLS402(Aman et al., 1992), DL221(de Graaff et al., 2016)), in contrast to fusion-negative liposarcoma cells (SW872) (Fig. 1). SNF2H and BRG1 PLA foci had relatively stronger signal compared to CHD4 PLA foci (Fig. 1, A and B). To confirm these interactions we performed co-immunoprecipitations (co-IPs) in the three human sarcoma cell lines (Fig. 1C). MLS402 expresses the 7-2 translocation FUS-CHOP variant (approx. 75 kDa); MLS1765 expresses the 13-2 translocation FUS-CHOP variant (approx. 100 kDa); and SW872 (negative control)(Stratford et al., 2012). We detected an interaction between FUS-CHOP and BRG1 as previously reported(Lindén et al., 2019). We also discovered a new interaction between FUS-CHOP and SNF2H of the ISWI complex (Fig. 1, C and D). CHD4 did not interact with FUS-CHOP in either fusion-expressing cell line (Fig. 1E). The controls ACF1, BAF170, and HDAC1 are known interactors with the chromatin remodelers ISWI, BAF, and NuRD, respectively (Fig. 1).

**Fig. 1.**
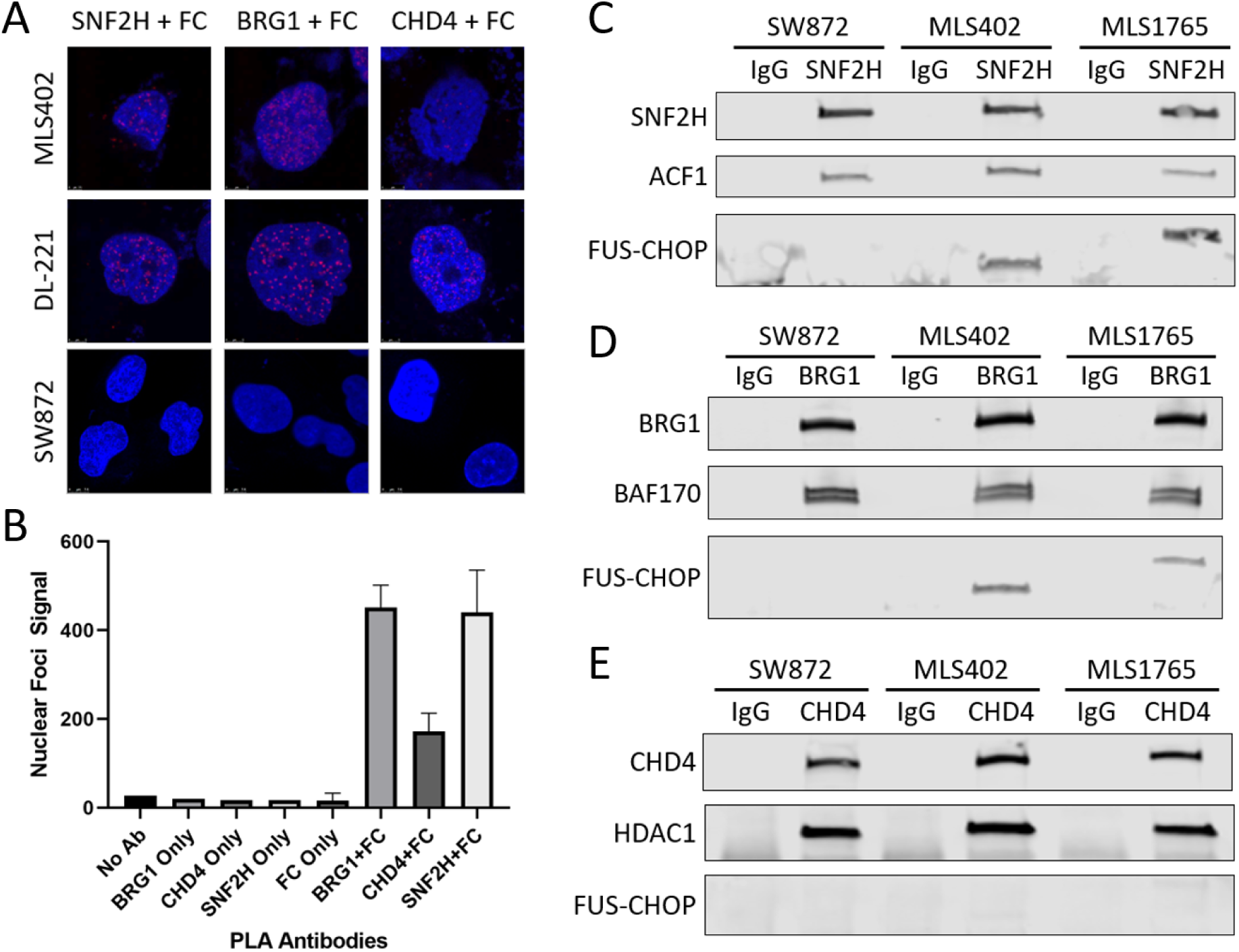
FUS-CHOP interacts with multiple chromatin remodeling complexes. (A) Confocal images of fusion-positive (DL-221, MLS402) and fusion-negative (SW872) nuclei stained with DAPI (blue). Red foci mark proteins that are in close proximity to FUS-CHOP (FC). SNF2H, BRG1, and CHD4 are ATPase subunits of their respective chromatin remodeling complexes. (B) Quantification of red foci in DL-221 nuclei with either single or paired PLA antibodies. (C) SNF2H co-IP for FUS-CHOP and the ACF1 subunit of ISWI complexes. (D) BRG1 co-IP for FUS-CHOP and the BAF170 subunit of the BAF complex. (E) CHD4 co-IP for FUS-CHOP and the HDAC1 subunit of the NuRD complex.

To further validate these interactions in a different organism, we performed co-IPs in mouse cell lines and mouse tumor cell lines with and without FUS-CHOP (fig. S1)(Chen et al., 2019; Kirsch et al., 2007). The FUS-CHOP-expressing tumor cell lines used for these experiments were derived from the conditional mouse model of FUS-CHOP-driven sarcoma that we generated using Cre and CRISPR/Cas9 technology(Chen et al., 2019). FUS-CHOP co-immunoprecipitated with Snf2h and Brg1 in these mouse cell lines (fig. S1). As observed in the human tumor cells, FUS-CHOP was not detected in the Chd4 co-IP. Therefore, the mouse co-IP validated the results of the human co-IP in support of this interaction in multiple different cell lines and across species.

### FUS-CHOP-driven sarcoma cells are dependent on SNF2H and BRG1 for proliferation

To determine if SNF2H and BRG1 impact the proliferation of FUS-CHOP-expressing cells, we stably knocked down Snf2h and Brg1 in KP (KrasG12D; p53-/-)(Kirsch et al., 2007) and 1650 (FUS-CHOP positive)(Chen et al., 2019) mouse sarcoma cells (Fig. 2A). Silencing Snf2h or Brg1 decreased proliferation of 1650 FUS-CHOP-driven mouse sarcoma cells (Fig. 2B). However, proliferation was unaffected in KP murine sarcoma cell lines. These data indicate that Snf2h and Brg1 are selectively important for the proliferation of FUS-CHOP-driven murine sarcomas.

**Fig. 2.**
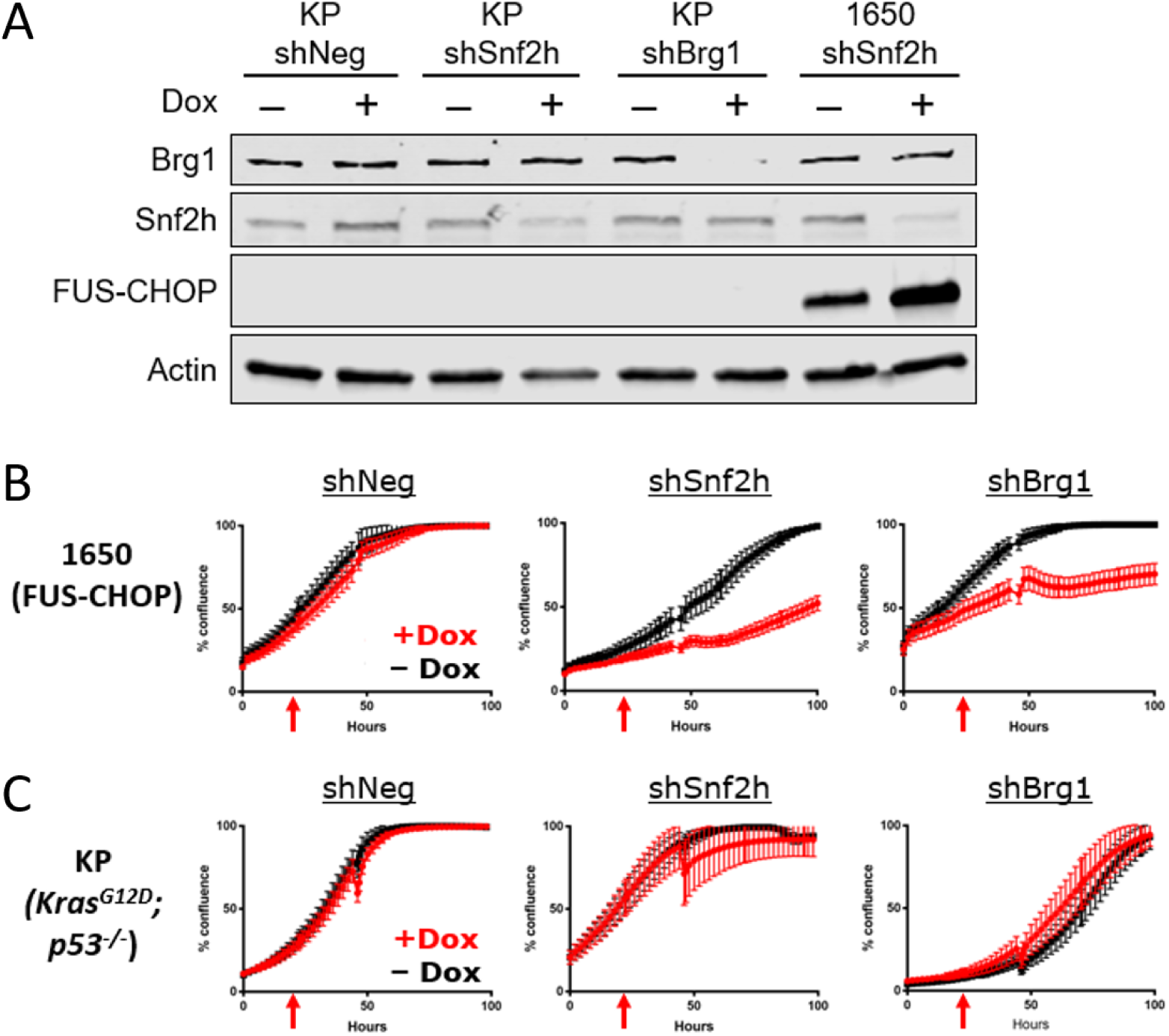
Snf2h and Brg1 are required for proliferation of FUS-CHOP-driven tumor cells. (A) Knockdown of Snf2h and Brg1 protein expression in 1650 (fusion-positive) and KP (fusion-negative) murine sarcoma cell lines. (B) Effect of Snf2h and Brg1 knockdown on proliferation of FUS-CHOP-driven murine sarcoma cell lines and (C) KP (fusion-negative) murine sarcoma cell lines. Red arrows denote addition of doxycycline (Dox) 24 hours after plating to induce shRNAs.

We then explored the stability of the interaction between SNF2H and FUS-CHOP and its dependence on nucleic acids (fig. S1). Interactions between SNF2H and FUS-CHOP were less stable than those between ISWI complex members SNF2H and ACF1 (fig. S1, D and E). The decreased stability of FUS-CHOP with the ISWI complex suggests a transient interaction, similar to EWS-FLI1 interaction with the BAF complex (Boulay et al., 2017). Although FUS-CHOP interacts with SNF2H independently of RNA or DNA, the interaction seems to be enhanced in the absence of RNA suggesting that RNA may modulate this interaction (fig. S1, F and G)(Maharana et al., 2018).

### FUS-CHOP retargets SNF2H to enhancers regulating proliferation

Because of the importance of the relationship between FUS-CHOP and SNF2H, we investigated the mechanism by which FUS-CHOP and SNF2H contribute to sarcomagenesis in MLPS. We first explored the genomic targeting of FUS-CHOP. To map FUS-CHOP binding sites genome-wide, we performed ChIP-seq in human MLPS cells as well as the FUS-CHOP-negative human liposarcoma cell line as a control (Fig. 3, fig S2). Distinct FUS-CHOP binding sites were detected in fusion-positive, but not fusion-negative chromatin (Fig. 3A, fig. S2A). 14,146 and 11,395 FUS-CHOP binding sites were identified in the MLS402 and DL221 cell lines, respectively (fig. S2. B and C). To increase the stringency of the analysis, we focused on those shared 5399 FUS-CHOP-bound sites detected in both MLS402 and DL221 cells. The FUS-CHOP binding sites were primarily located in intronic and distal intergenic regions with 5,197 FUS-CHOP binding sites at enhancers (fig. S2D). Since EWS-FLI1 is known to target GGAA repeats, we searched for repetitive sequence motifs, but no such repeat sequences were identified through our analysis. To determine whether SNF2H colocalized with FUS-CHOP on chromatin, we performed SNF2H CUT&RUN (Fig. 3, A and C, fig. S2F-H). We identified 8,344 sites of SNF2H binding that were shared by the FUS-CHOP positive MLPS cell lines (but not in the fusion-negative cells) (Fig. 3B, fig. S2, G and H). Of these sites, we identified about 10% that were also bound by FUS-CHOP (Fig. 3B). H3K27 acetylation at the sites targeted by FUS-CHOP and SNF2H in MLS402 and DL221 cell lines indicates that they represent active enhancer elements (Fig. 3C-E). Interestingly, modest H3K27ac in the FUS-CHOP-negative SW872 cells suggests that other transcription factors may mediate acetylation at some of these sites. FUS-CHOP bound at these enhancers correlates with the abundance of SNF2H and H3K27ac and supports a model of recruitment of SNF2H to these enhancers defined by FUS-CHOP. DNA motif analysis of shared FUS-CHOP and SNF2H binding sites revealed significant enrichment of FOSL1, REL, and RUNX motifs, relative to genomic background (Fig. 3I). Interestingly, despite detection of the DDIT3 (CHOP) motif, it was less enriched than FOSL1. SNF2H/FUS-CHOP shared sites demonstrate similarly enriched DNA-binding motifs as those bound by FUS-CHOP (Fig. 3I, fig. S2E). Interestingly, the DDIT3 (or CHOP) motif was a prominent shared motif, but not a prominent motif in FUS-CHOP binding alone suggesting that some less prevalent motifs (such as DDIT3) are enriched at shared FUS-CHOP and SNF2H binding sites (fig. S2E). Wild type CHOP is implicated in the unfolded protein response (UPR), and under stress conditions, CHOP and ATF4 bind to very similar sequences(Han et al., 2013). However, we observed a different set of DNA-binding motifs in our analysis. Differences in DNA binding between FUS-CHOP and wild type CHOP may explain how the fusion protein is oncogenic while the wild type CHOP is pro-apoptotic and initiates ER stress-induced cell death(Barone et al., 1994). Yet, the binding of FUS-CHOP to “classical” DDIT3 sequence motifs also supports a model where FUS-CHOP displaces or prevents wild type CHOP from binding its transcriptional targets important for growth arrest or apoptosis. Further experiments are required to elucidate if this “dominant negative” effect occurs either at the DNA/chromatin level or through protein-protein interactions, though evidence in the literature suggests the former mechanism is more likely(Barone et al., 1994; Crozat et al., 1993; Zinszner et al., 1994).

**Fig. 3.**
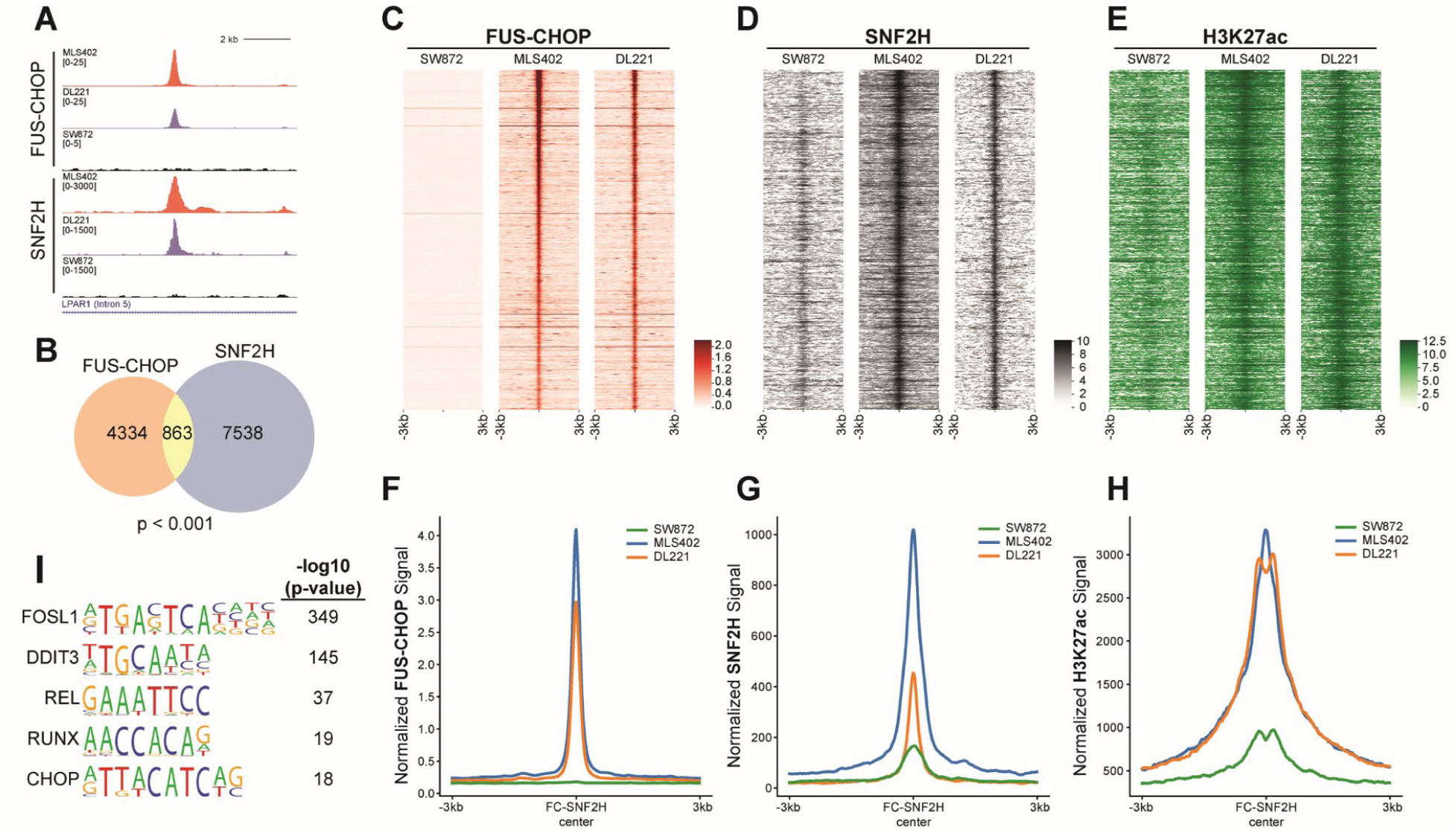
FUS-CHOP colocalizes with SNF2H at active enhancers. (A) Representative tracks of a FUS-CHOP SNF2H shared binding site. (B) Venn diagram showing the overlap of FUS-CHOP ROE at enhancers and SNF2H ROE (Permutation test, p < 0.001). (C-E) Heatmaps of FUS-CHOP, SNF2H, and H3K27ac signal densities in human fusion-positive (DL221, MLS402) and fusion-negative (SW872) liposarcoma cell lines. 6-kb windows in each panel are centered on FUS-CHOP SNF2H shared binding sites (n = 863), ranked by significance of overlapping peak calls. (F-H) Average line plots of FUS-CHOP, SNF2H, and H3K27ac signal at ROE bound by both FUS-CHOP and SNF2H. SNF2H signal is increased at these enhancers in the presence of FUS-CHOP. H3K27ac signal is also increased at these enhancers in the presence of FUS-CHOP and SNF2H. The x-axis represents a 6-kb window centered on FUS-CHOP SNF2H binding sites. (I) Enriched motifs found at overlapping FUS-CHOP and SNF2H regions of enrichment.

To determine how the shared occupancy of SNF2H and FUS-CHOP impacted gene expression, we performed RNA-seq on human MLPS and liposarcoma cell lines and associated the transcriptomic differences between these cell lines with regions of shared occupancy (fig. S3). Our analysis revealed that regions of shared FUS-CHOP and SNF2H enrichment were associated with 711 genes, of which 338 genes were upregulated and 83 genes were downregulated in the DL221 cell line relative to SW872 (fig. S3A). These regions were associated with 212 upregulated and 144 downregulated genes in the MLS402 MLPS cell line compared to SW872 (fig. S3B). There was a strong correlation between the magnitude of both gene activation and repression in DL221 and MLS402 cell lines at these genes (fig. S3, C and D). These data show that the enhancers occupied by FUS-CHOP and SNF2H upregulate a common set of genes. Activated genes are enriched for those involved in cell proliferation, suggesting a direct role for that FUS-CHOP function in sarcomagenesis (fig. S3, E and F) (Chen et al., 2013; Kuleshov et al., 2016).

### The FUS-CHOP PrLD is phosphorylated after X-ray irradiation

The FUS-CHOP PrLD is required for transformation of NIH-3T3 cells suggesting that the PrLD may mediate transactivation in the fusion protein. Substitution of the PrLD by other transactivation domains does not strongly transform cells (Zinszner et al., 1994). In contrast, substitution of FUS with EWS in the fusion oncoprotein results in similar transformation(Zinszner et al., 1994) and the EWS-CHOP fusion is also associated with MLPS in patients(Dal Cin et al., 1997; Panagopoulos et al., 1996). These results indicate that FUS and EWS confer specific activities necessary for transformation. To investigate a potential link between the role of FUS PrLD as a transactivator and the exquisite clinical response of MLPS to radiotherapy, we hypothesized that the interaction of FUS-CHOP with binding partners, such as SNF2H, might be regulated through DNA damage-induced phosphorylation of the FUS-CHOP PrLD, which could interfere with chromatin remodeler interactions and contribute to the sensitivity of MLPS to ionizing radiation.

Immunoblotting with antibodies that recognize phospho-SQ/TQ (pSQ/TQ) sites, we explored the phosphorylation state of FUS-CHOP immunoprecipitated from cells treated with ATM and DNA-PK inhibitors. We found that FUS-CHOP was phosphorylated after irradiation by DNA-PK and ATM (fig. S4). pSQ/TQ phosphorylation levels following irradiation in the presence of the DNA-PK inhibitor NU-7026 were similar to unirradiated cells (fig. S4B). Using a human FUS phospho-Ser42 specific antibody, we observed that sites in the PrLD were phosphorylated in FUS-CHOP driven mouse sarcoma cells (fig. S4C). These data show that, like wild type FUS, FUS-CHOP is phosphorylated by DNA-PK and ATM after irradiation(Gardiner et al., 2008).

### PrLD phosphorylation inhibits interactions with SNF2H and transformation

Next, we investigated if radiation affects the interaction of FUS-CHOP with SNF2H. 45 minutes after 10 Gy X-ray irradiation, we observed a significant decrease in the FUS-CHOP interaction with SNF2H (Fig. 4A). To directly test if the decreased interaction of FUS-CHOP with SNF2H was due to phosphorylation of the PrLD, we generated wild type (FCWT), phospho-dead (FC-12A) or phospho-mimic (FC-12E) FUS-CHOP mutants at the twelve SQ/TQ sites in the FUS-CHOP PrLD. These FUS-CHOP variants were stably expressed in NIH-3T3 cells (Fig. 4, B and C)(Monahan et al., 2017). In contrast to the wild type FUS-CHOP and the phospho-dead 12A mutant, SNF2H interaction with the phospho-mimic 12E mutant was diminished (Fig. 4D). We then tested the transformation activity of these mutants by soft agar colony formation. Wild type and the 12A mutant, but not the 12E mutant, transformed NIH-3T3 cells (Fig. 4E). These data suggest that phosphorylation of the FUS-CHOP PrLD disrupts functions of the fusion protein necessary for transformation and proliferation, including interactions with SNF2H and potentially other nuclear proteins.

**Fig. 4.**
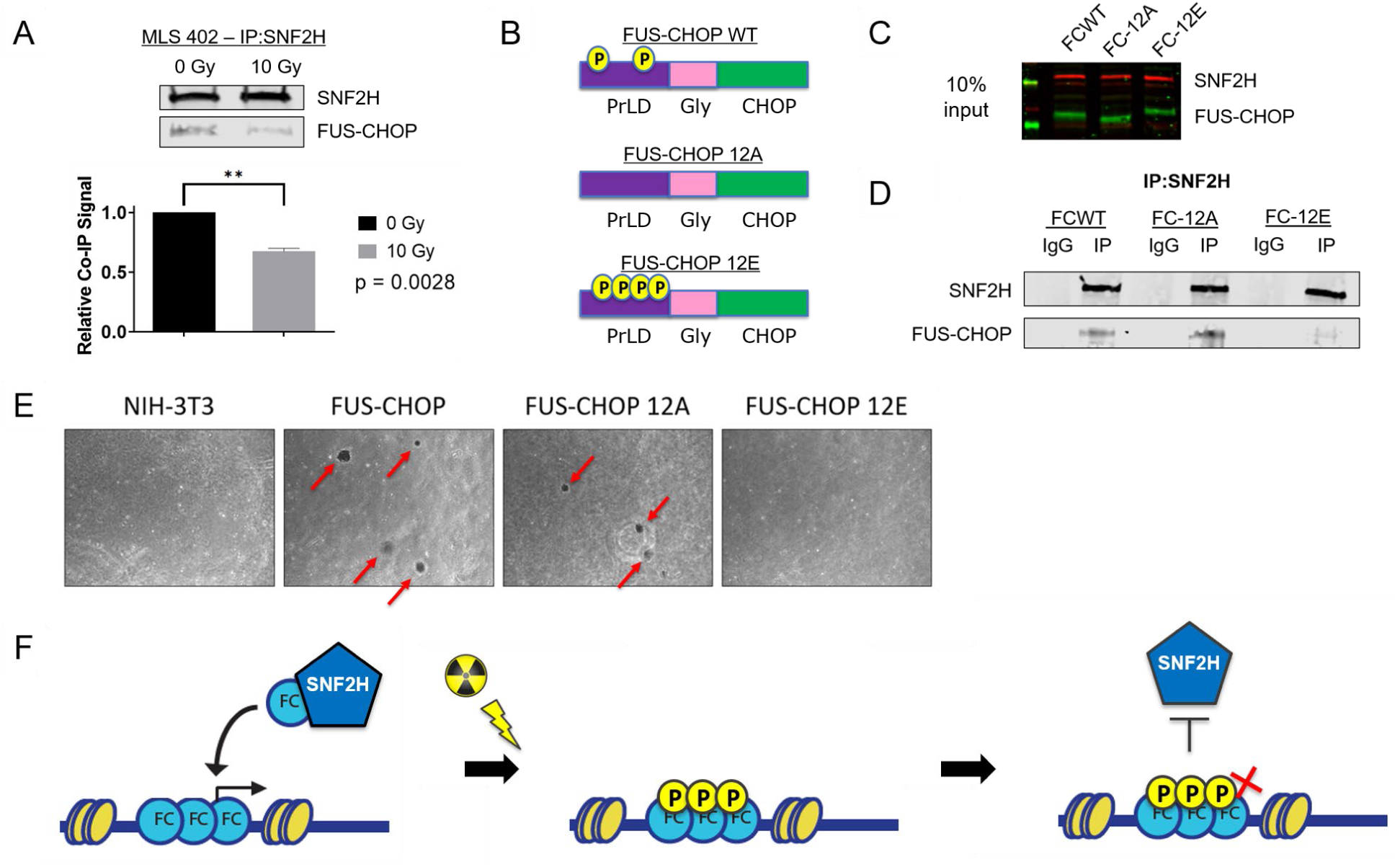
Ionizing radiation decreases protein-protein interactions between SNF2H and FUS-CHOP by DNA damage-induced phosphorylation of the PrLD. (A) Co-IP of SNF2H and FUS-CHOP 45 minutes after either sham or 10 Gy X-ray irradiation. ** p = 0.0028, unpaired t-test. (B) Schematic of FUS-CHOP wild type, phospho-dead (12A), and phospho-mimic (12E) cDNA constructs. (C) NIH-3T3 cells stably expressing FUS-CHOP and its phosphomutant forms. (D) Co-IP of SNF2H in NIH-3T3 cell lines stably expressing FUS-CHOP (FCWT), FUS-CHOP 12A (FC-12A), and FUS-CHOP 12E (FC-12E). (E) Soft agar colony formation assay to evaluate cellular transformation using NIH-3T3 cells that stably express either wild type FUS-CHOP or phosphomutant forms of FUS-CHOP. (F) Model of FUS-CHOP mediated oncogenic activity and radiosensitivity in MLPS. FUS-CHOP and SNF2H interact to target enhancers and activate genes, but after irradiation FUS-CHOP is phosphorylated in the PrLD, which interferes with SNF2H interactions and MLPS maintenance.

A recently appreciated property of proteins containing PrLDs, including the FET proteins, is the ability to form phase-separated compartments, or biomolecular condensates. Indeed, it is possible that FUS-CHOP activates enhancers and gene targets by forming nuclear condensates that contain chromatin remodelers and transcriptional machinery. We demonstrated that specific protein-protein interactions between the PrLD of FUS-CHOP and SNF2H can be disrupted by using a phospho-mimic mutant of FUS-CHOP (Fig. 4D). This decreased interaction between FUS-CHOP and SNF2H following irradiation may result from increased solubility of the phosphorylated form of FUS-CHOP mediated by DNA damage. Using purified proteins, we observed that the FUS-CHOP 12E phosphomimetic mutant remains soluble after cleavage of an MBP solubility tag (fig. S4D). In contrast, wild type and the 12A mutant proteins aggregate into micron-scale particles.

## Discussion

Our work demonstrated that retargeting chromatin remodeling complexes by FET fusion proteins may be a more broadly applicable mechanism beyond EWS-FLI1 in Ewing sarcomagenesis (Boulay et al., 2017). Importantly, we showed that in addition to the BAF complex, FET proteins can interact with other chromatin remodeling complexes such as the ISWI complex, which contains the SNF2H ATPase subunit (Fig. 1, fig. S1). We also showed that the SNF2H and BRG1 ATPase subunits were important for proliferation of FUS-CHOP-driven sarcomas (Fig. 2). We hypothesized that proliferation of FUS-CHOP-driven sarcomas may be dependent on these chromatin remodelers because recruitment of chromatin remodeling complexes to DNA regulatory elements by FUS-CHOP was important for sarcomagenesis. To test this model of oncogenesis, we first mapped FUS-CHOP and SNF2H binding sites genome-wide (Fig. 3, fig S2). These two proteins were found to co-localize at active enhancers in human MLPS cell lines that upregulated a common set of genes implicated in cell proliferation, migration, apoptosis, and several other processes (Fig. S3). This general mechanism of FET fusion-mediated chromatin remodeling may be a model for sarcomagenesis of an entire class of translocation-positive sarcomas that contain FET protein translocation partners. Additionally, to address lessons from modeling sarcomas in other organisms, this mechanism may explain why fusion-positive sarcomas are difficult to model in mice where the specific DNA-binding motifs and enhancers required for sarcomagenesis in humans differ from those required for these processes in mice(Chen et al., 2019; Minas et al., 2016).

An intriguing and emerging property of proteins containing PrLDs, such as FET proteins, is the ability to form phase-separated biomolecular condensates(Banani et al., 2017; Maharana et al., 2018; Ryan et al., 2019; Shin et al., 2017). As such, FET proteins can phase separate under physiological conditions, and perhaps most interestingly, these intrinsically disordered regions (IDRs) can be post-translationally modified to promote or prevent phase separation and other intermolecular interactions. Given that the IDR of FUS is translocated in FUS-CHOP, we began to view “retargeting of chromatin remodelers” through the lens of biomolecular condensates and phase separation. Indeed, FUS-CHOP may be forming intranuclear condensates that contain chromatin remodelers and transcriptional machinery to activate enhancers and gene targets. Although our work is limited to the level of chromatin and does not directly explore formation of biomolecular condensates, we demonstrate specific protein-protein interactions between the PrLD of FUS-CHOP and SNF2H can be disrupted by using a phospho-mimic mutant of FUS-CHOP (Fig. 4). Post-translational modifications of FET PrLDs can change the charge of these proteins and either promote or prevent phase separation. In the case of FUS, phosphorylation of the domain at twelve phospho-SQ/TQ sites in the PrLD has been shown to increase the solubility of FUS and to disrupt phase separation (Monahan et al., 2017; Rhoads et al., 2018). Taken together, our results suggest a model linking oncogenic activity and radiosensitivity in MLPS supporting a critical role for the PrLD of FUS-CHOP to mediate oncogenic retargeting of SNF2H (Fig. 4F). Phosphorylation of the PrLD by activated DNA damage response kinases alters the solubility of FUS-CHOP, which destabilizes FUS-CHOP/SNF2H interactions necessary for MLPS maintenance. Therefore, the clinically relevant radiosensitivity of MLPS may be mediated by radiation-induced disruption of the phase separation properties of FUS-CHOP and diminishing protein-protein interactions required for tumor maintenance. Drugs that promote phosphorylation of the PrLD in FET proteins like FUS that disrupt interactions with chromatin remodeling complexes may be effective treatments for FET fusion positive sarcomas.

## Acknowledgements

We thank Pierre Åman (University of Gothenberg) for cell lines MLS402 and MLS1765, and thank Keila Torres and Alexander Lazar (MD Anderson) for cell line DL221. We thank Steven Henikoff, Derek Janssens, and Nan Liu for their advice, shared reagents, and protocols for CUT&RUN. We also thank Andrea Ventura (Memorial Sloan Kettering) for helpful suggestions and discussion. This work was supported by the National Cancer Institute of the US, NIH (awards R35 CA197616 to DGK, F30 CA206424 to MC) and the T32 GM007171 MSTP training grant (Duke University).

## Author contributions

MC and DGK contributed to the original project conceptualization, methodology, supervision, administration, funding, resources, visualization, original writing, and review and editing of the manuscript. JPF and IJD contributed to methodology, resources, investigation, visualization, data curation, and formal analysis. MC, JPF, ICL, NHL, ARD, WF, and ESX contributed to investigation, resources, validation, formal analysis, and review and editing of the manuscript.

## Data and materials availability

All data generated or analyzed in this study are included in this published article and its supplementary materials. All genomic data underlying the study are available/will be deposited at https://www.ncbi.nlm.nih.gov/geo/query/acc.cgi?acc=GSE145678 Access will be made public after publication.

## STAR Methods

### RESOURCE AVAILABILITY

#### Lead Contact

Further information and requests for resources and reagents should be directed to and will be fulfilled by the Lead Contact, David Kirsch.

#### Materials Availability

Plasmids generated in this study will be deposited to Addgene or are available through Addgene.

#### Data and Code Availability

All genomic data underlying the study are available/will be deposited at https://www.ncbi.nlm.nih.gov/geo/query/acc.cgi?acc=GSE145678 Access will be made public after publication.

### EXPERIMENTAL MODEL DETAILS

#### Tissue culture and cell line generation

NIH-3T3 cell lines were purchased from ATCC (CRL-1658) and cultured in DMEM media (ThermoFisher Scientific, 11965092) supplemented with 10% fetal bovine serum (ThermoFisher Scientific, 16000044) and 1% antibiotic–antimycotic (ThermoFisher Scientific, 10091148), and incubated at 37°C with 5% CO2 in a humidified cell-culture incubator. Tumor cells were isolated from primary mouse sarcomas as previously described(Chen et al., 2019). Briefly, tumor tissue was minced in the cell-culture hood and digested by dissociation buffer in PBS (ThermoFisher Scientific, 14040133) containing collagenase type IV (5 mg/ml, ThermoFisher Scientific, 17104-019), dispase (1.3 mg/ml, ThermoFisher Scientific, 17105-041) as well as trypsin (0.05%, ThermoFisher Scientific, 25200056) for about 1 h at 37°C. Cells were washed with PBS (ThermoFisher Scientific, 10010023) and filtered using a 40 mm sieve (Corning, 431750), and cultured for at least four passages to deplete stroma before being used for experiments. The SW872 cell line was purchased from ATCC. MLPS cell lines MLS402 and MLS1765 were gifts from Pierre Åman (University of Gothenburg, Sweden), and the DL-221 cell line was a gift from Keila Torres and Alexander Lazar (University of Texas MD Anderson Cancer Center, Houston, Texas). SW872, MLS402 and MLS1765 were cultured in RPMI with 10% FBS and 1% antibiotic-antimycotic. Inducible shRNA cell lines were cultures in DMEM with 10% Tet-free FBS (Takara) and 1% antibiotic-antimycotic. DL-221 and all other cell lines were cultured in DMEM with 10% FBS and 1% antibiotic-antimycotic.

### METHOD DETAILS

#### Plasmids and Vectors

FUS-CHOP WT, FUS-CHOP 12A, and FUS-CHOP 12E constructs were designed in Snapgene and synthesized by Genscript. The cDNA fragments were then subcloned into N174-MCS (Puro) vector backbone, which was a gift from Adam Karpf (Addgene plasmid # 81068 ; http://n2t.net/addgene:81068 ; RRID:Addgene_81068). shRNA sequences from the RNAi consortium were obtained from the Sigma-Aldrich shRNA database and shRNA oligos were ordered from Integrated DNA Technologies (IDT). shRNAs were subcloned into Tet-pLKO-puro, which was a gift from Dmitri Wiederschain (Addgene plasmid # 21915 ; http://n2t.net/addgene:21915 ; RRID:Addgene_21915) (Wiederschain et al., 2009).

#### Radiation treatments

Cells were cultured at least 24 hours prior to irradiation experiments. The X-RAD 160 (Precision X-Ray) cell irradiation system was used at 160 kVp x 18 mA energy. An F1 filter (2 mm aluminum) was used for beam conditioning. Sample distance was set to 40 cm. For a dose of 10 Gy, calculations based on annual dosimetry determined a treatment duration of 230 seconds.

#### Immunoblotting and antibodies

Samples were lysed in RIPA buffer (50 mM Tris-HCl, pH 8.0 with 150 mM NaCl, 1% NP-40, 0.5% sodium deoxycholate, 0.1% sodium dodecyl sulfate (SDS)) for 30 min on ice (Sigma-Aldrich, R0278), sonicated briefly, then centrifuged at 10,000x g for 20 min at 4°C. Protein concentration was determined for the lysate supernatant by BCA assay (Pierce, 23225). Samples was boiled in 4X Laemmli sample buffer (Bio-Rad, 1610747) at 95°C for 5 min, then cooled to room temperature before loading in a 4-20% Tris-Bis polyacrylamide gel. Samples were electrophoresed at 200 V for 30 min before transfer to nitrocellulose. Membranes were blocked in 5% non-fat dry milk or 5% bovine serum albumin (BSA) in Tris-buffered saline (TBS, Corning, 46-012-CM). Next, membranes were incubated overnight at 4°C with primary antibodies diluted in TBS-T (0.1% Tween-20) with 3% BSA: CHOP 1:1,000 (Cell Signaling Technology, 2895S); CHD4 1:1000 (D8B12, Cell Signaling Technology); BRG1 1:1000 (D1Q7F, Cell Signaling Technology); SNF2H 1:5000 (ABE1026, EMD Millipore); BAF170 1:1000 (D8O9V, Cell Signaling Technology); ACF1 1:1000 (ab187670, abcam); HDAC1 1:1000 (10E2, Cell Signaling Technology); TBP 1:1000 (D5C9H, Cell Signaling Technology). Membranes were washed three times in TBS-T for 5 min before secondary antibody incubation with goat anti-rabbit IRDye800 (Li-Cor Biosciences, P/N 925-32211) and goat anti-mouse IRDye680 (Li-Cor Biosciences, P/N 925-68070) both at 1:10,000 dilutions in TBS-T for 1 h at room temperature. The membranes were washed three times in TBS-T for 5 min and imaged using an Odyssey CLx (Li-Cor Biosciences). Image analysis for normalization and quantification were performed using Image Studio (Version 5.2, Li-Cor Biosciences, P/N 9140-500).

#### Co-immunoprecipitation

The Co-IP protocol was adapted from Boulay et al.(Boulay et al., 2017). Briefly, media was aspirated and washed twice with cold PBS. On ice, 1 mL of cytoplasmic lysis buffer (Buffer A: 10 mM Tris-HCl pH 8.0, 10 mM KCl, 1.5 mM MgCl2, 0.15% NP-40, 1 mM PMSF, 1 mM DTT, 1 tablet PhosSTOP and 1 tablet cOmplete protease inhibitor per 10 mL buffer) was added per 10 cm dish and scraped into a tube. Cells were incubated on ice for 5 minutes in Buffer A with periodic pipetting to mix and shear cells. Nuclei were pelleted at 1200 rpm for 5 minutes, and supernatant was aspirated carefully. The nuclei were washed with Buffer A without NP-40 and pelleted again via centrifugation at 1000 x g for 5 minutes. The supernatant was aspirated and nuclei were resuspended in nuclear lysis buffer (Buffer B: 50 mM Tris-HCl pH 8.0, 150 mM NaCl, 5 mM EDTA, 0.5% NP-40, 10% glycerol, 1 mM PMSF, 1 tablet PhosSTOP and 1 tablet cOmplete protease inhibitor per 10 mL buffer), approximately 5-6 volumes for adequate lysis. Gentle sonication with a QSONICA 800 R instrument at 20 Amp, 1 sec on/1 sec off, for 60 seconds was used to help lyse nuclei. After incubation for 30 minutes, nuclear lysate was centrifuged for 15 minutes at 14000 rpm at 4°C. The nuclear fraction was collected and quantified, then the final concentration was adjusted to 1 mg/mL with Buffer B. Each IP fraction was incubated with 3 ug of antibody (IgG rabbit, Cell Signaling Technology; CHD4 (D8B12), Cell Signaling Technology; BRG1 (D1Q7F), Cell Signaling Technology; SNF2H (ABE1026), EMD Millipore) or (1:50 dilution CHOP (L63F7), Cell Signaling Technology) overnight at 4°C on a tube rotator. Dynabeads were resuspended in the vial by vortexing >30 seconds and enough beads for 20 uL of beads for each IP reaction were transferred to a new tube. The tube was placed on a magnet to separate the beads from the solution, and the beads were washed once with Buffer B. The IP samples were then incubated with 20 uL of protein G Dynabeads for 2 hours at 4°C on the tube rotator. Samples were then washed 5 times at 4°C with 500 uL of Buffer B with gentle pipetting or inversion on a magnetic rack. After the final wash, the beads were resuspended in 30 uL 2x gel loading buffer with DTT, boiled, and processed using a standard western blotting protocol.

#### Urea denaturation assay

Urea denaturation was performed exactly as the co-IP protocol above with one exception. Before incubation with the primary antibody overnight, lysate was incubated at room temperature for 15 minutes with various concentration of urea (0M, 0.5M, 1M, 2M).

#### RNase/DNase assay

RNase/DNase assays were performed exactly as the co-IP protocol above with two changes. Before incubation with the primary antibody overnight, lysate was incubated at 37°C for 15 minutes with RNase A in Buffer B at 100 ug/mL then incubated with IP antibody overnight at 4°C for the RNase assay. Sample lysate was incubated at 37°C for 15 minutes with 100 units/mL DNase in Buffer B then incubated with IP antibody overnight at 4°C.

#### Soft agar transformation assays

1.8% Bactoagar was made with diH2O and autoclaved. 2x and 1x DMEM were prepared using DMEM powder (ThermoFisher Scientific, 12100046) fetal bovine serum (ThermoFisher Scientific, 16000044), and antibiotic-antimycotic (ThermoFisher Scientific, 10091148). 0.6% agar was made by diluting 1.8% agar with DMEM and kept in a 42°C water bath. 3 mL of 0.6% agar was poured per well of 6-well plates and allowed to solidify in a hood for 10 minutes before transferring to an incubator. Cells were trypsinized and counted using Trypan Blue solution (ThermoFisher Scientific, 15250061). 250 μL of cells and 500 μL of 0.6% agar were mixed and gently pipetted onto the bottom agar in each well to create the 0.4% agar top layer. 5000 cells were plated per well in triplicate for each cell line. After plating, plates were placed in an incubator and allowed to grow for 3-4 weeks. Media was supplemented each week by adding 0.5 mL of DMEM per well to prevent drying.

#### Proximity ligation assay (PLA)

The Duolink PLA red mouse/rabbit kit was used to perform proximity ligation assay per manufacturer’s protocols. Briefly, cells were plated in 8-well chamber slides overnight. The next day, media was aspirated, cells were fixed with 4% formaldehyde with methanol permeabilization. Cells were then blocked with Duolink blocking solution and incubated with primary antibody overnight. Next, Duolink PLA probes were incubated with the samples followed by ligation and amplification. Finally, the slides were mounted with a coverslip using Duolink in situ mounting medium with DAPI, and the slides were either imaged 15 minutes later or kept at -20°C until imaging using a Leica SP5 Inverted Confocal Microscope (DMI6000CS) with the 405 nm and 647 nm lasers for excitation.

#### ChIP-seq

Chromatin was prepared using a SimpleChIP plus Sonication Chromatin IP Kit from Cell Signaling Technology according to manufacturer’s instructions following optimization of crosslinking, sonication and antibody conditions for each cell line. Briefly, one chromatin preparation for each cell line was prepared from approximately 20 million cells. Cells were crosslinked in 1% formaldehyde (methanol-free) for 9 minutes at room temperature. Glycine neutralization followed for 5 minutes at room temperature followed by 2x wash with cold PBS. Next, cells were scraped into cold 1x PBS with protease inhibitor, then pelleted at 1000 x g for 5 minutes at 4C. PBS was removed and cells were resuspended in 1X cell lysis buffer with protease inhibitor for 10 minutes prior to nuclei preparation. Nuclei were pelleted at 5000 x g for 5 minutes at 4C and the pellet was resuspended in 1X cell lysis buffer with protease inhibitor for 5 minutes before pelleting at 5000 x g for 5 minutes at 4C. Pelleted nuclei were then resuspended in cold nuclear lysis buffer with protease inhibitor and incubated on ice for 10 minutes. 1 mL of the nuclear lysate was transferred to appropriate tubes for sonication. Chromatin was sonicated using a Qsonica Q700 sonicator with Cup Horn and chiller (70 Amp, 15 sec on, 45 sec off) for the following durations per cell line: SW872, 15 minutes; DL221, 12 minutes; MLS402, 20 minutes. Following sonication, lysates were clarified by centrifugation at 21000 x g for 10 minutes at 4C and supernatant was transferred to a new tube. 50 uL of each lysate was taken for analysis of fragmentation and concentration determination. 10 ug ChIP DNA was used per IP sample. Samples were incubated with primary antibody overnight at 4C on a tube rotator. The next day, samples were incubated with protein G magnetic beads for 2 hours at 4C. Beads were isolated and washed 3 times in low salt wash buffer and 1 time in high salt wash buffer. Finally, chromatin was eluted from magnetic beads, protein was reverse crosslinked, and DNA was purified and pooled for library prep and sequencing.

Adaptor sequences were removed from reads with cutadapt (v.1.12). FASTX-Toolkit (v0.0.12) with options -Q 33, -p 90, and q 20 was used to filter reads. Reads were aligned to the hg19 genome using STAR (v2.5.2b) with the following options: --outFilterMismatchNmax 2, -- chimSegmentMin 15, --chimJunctionOverhangMin 15, --outSAMtype BAM Unsorted, -- outFilterType BySJout, --outFilterScoreMin 1, --outFilterMultimapNmax 1. Tracks of FUS-CHOP were created by visualizing RPM normalized bigWigs on UCSC genome browser. FUS-CHOP regions of enrichment (ROE) were identified using MACS2 (v2.1.2) using default parameters and cell line matched input samples as controls. For each ROE identified by MACS2 a score per million (SPM) was calculated, and low confidence peaks (bottom 5% of SPM scores) were removed for downstream analysis(Corces et al., 2018). FUS-CHOP ROE was defined as +/- 90 bp MACS2 identified summit. The +/- 90 bp window size was determined empirically by calculating the ratio of the number of overlapping FUS-CHOP ROE summits found in the MLS402 and DL221 cell line over the total number of MLS402 ROE as a variable window was added to ROE summits found in the MLS402 cell line. A 180 bp window was selected for downstream analysis as an increase in the window size did not increase the overlapping number of ROE. Annotating FUS-CHOP ROE with genomic features was done using the HOMER annotatePeaks.pl with the options hg19 (v4.10.3). FUS-CHOP ROE-enhancer set was defined by excluding any FUS-CHOP ROE that fell with a gene promoter region (+300:Ref-seqTSS:-500 bp – longest transcript). Average line plot signal for FUS-CHOP and H3K27ac in MLS402, DL221, and SW872 cells was isolated using deepTools (v3.2.0). Signal shown in heatmaps was log2 transformed, and the max signal was set at the 99th percentile across MLS402, DL221, and SW872 cell lines. Motifs found at FUS-CHOP ROE were identified using HOMER (v4.10.3) using HOMER defined random background regions. GREAT (McLean et al., 2010) was used to annotate ChIP-seq ROE with associated genes. All visualizations were created using Python. Any ChIP-seq intersection set was identified using bedtools (v2.28.0).

### CUT&RUN

CUT&RUN was performed as published and as on protocols.io (Meers et al., 2019; Skene et al., 2018). Briefly, cells were bound to conacavalin beads, permeabilized with digitonin and incubated with primary antibody overnight. Protein A/G-MNase was used to digest chromatin for 30 minutes and soluble DNA was purified via phenol-chloroform extraction. DNA libraries were prepped and barcoded using NEBNext adapters and barcodes before pooling and sequencing at Genewiz.

Adaptor sequences were removed using (cutadapt v1.12), and reads were then quality filtered using fastq quality filter in FASTX-Toolkit (v0.0.12) with options -Q 33, -p 90, and q 20. Reads were aligned to a custom hg19-eColi-K12 genome using STAR (v.2.5.2b) with the following options: --outFilterMismatchNmax 2, --chimSegmentMin 15, -- chimJunctionOverhangMin 15, --outSAMtype BAM Unsorted, --outFilterType BySJout, -- alignMatesGapMax 1000, --outFilterScoreMin 1, --outFilterMultimapNmax 1. The following methods were applied to CUT&RUN experiments for SNF2H, and H3K27ac in MLS402, DL221, and SW872 cell lines. Tracks were created by using RPM normalized and spike-in adjusted bigWigs on UCSC genome browser. Regions of enrichment were identified using MACS2 (v2.1.2) using default parameters. ROE were annotated with a SPM score, and peaks with a score in the bottom 5% of SPM scores were not included in downstream analysis(Corces et al., 2018). The intersection of MSL402 SNF2H ROE and DL221 SNF2H was identified using bedtools (v2.28.0) with options -f 0.5 -r. Average line plot signal for SNF2H and H3K27ac was isolated using deepTools (v3.2.0). Heatmap signal shown was maxed out at 99th percentile across MLS402, DL221, and SW872 cell lines, and log2 transformed. The intersection of FUS-CHOP ChIP-seq ROE and SNF2H CUT&RUN ROE was identified using bedtools (v2.28.0). A permutation test was used to determine the significance of the FUS-CHOP SNF2H overlap. SNF2H ROE were held constant, while the FUS-CHOP ROE were permuted by identifying size and chromosome matched random regions excluding unmappable regions. The intersection for each permutation was determined with bedtools (v2.28.0) with options -f 0.5 -r. The number of overlapping regions were stored for each iteration, and the p-value was calculated as the total number of permutations with an intersection > 863 over the total number of permutations (n = 1000).

### RNA-seq

Cells were washed with PBS twice and lysed with TRIzol reagent. RNA was purified following manufacturer’s instructions using a DirectZol RNA miniprep kit (Zymo Research). RNA samples were spiked with ERCC for normalization and sent to Genewiz for sequencing. Adaptor sequences were trimmed using (cutadapt v1.12). Reads were then quality filtered using fastq quality filter in FASTX-Toolkit (v0.0.12) with options -Q 33, -p 90, and q 20. Reads were aligned to hg19 genome using STAR (v2.5.2b) with the following options: --quantMode TranscriptomeSAM, --outFilterMismatchNmax 2, --alignIntronMax 1000000, --alignIntronMin 20, --chimSegmentMin 15, --chimJunctionOverhangMin 15, --outSAMtype BAM Unsorted, -- outFilterType BySJout, --outFilterScoreMin 1. Gene expression (TPM) was created using Salmon (v0.11.3). ERCC spike-in was performed, but not used in downstream analysis. Differentially expressed genes (MLS402/DL221 v. SW872) were identified using DESeq2 (v1.24.0). MA plots were created using Python.

### Protein production and purification

Target sequences for MBP-FUS-CHOP WT, MBP-FUS-CHOP 12A, and MBP-FUS-CHOP 12E cDNA sequences were synthesized by Genscript and subcloned into an E. coli expression vector pET28a-MBP that contains 6xHis-MBPtag. A TEV sequence was inserted between the MBP tag and FUS-CHOP cDNAs for MBP tag release via cleavage by TEV protease. BL21 star (DE3) E. coli cells were transformed with the recombinant plasmids and a single colony was inoculated into Terrific Broth (TB) medium containing the appropriate antibiotic. A 2 L culture was incubated at 37°C and when the OD600 reached about 1.2, protein expression was induced with IPTG at 15C for 16 hours. Cells were harvested by centrifugation. Following centrifugation cell pellets were resuspended with lysis buffer followed by sonication. The supernatant was kept for further purification. The target proteins were purified via one-step purification on a Ni column followed by sterilization through a 0.22 µm filter. No visible precipitation was observed after purification and two cycles of freeze-thaw testing was performed to ensure no visible precipitation after freezing. pH 7.4 PBS with 10% glycerol was used as the final vehicle for the proteins. Proteins were analyzed via SDS-PAGE and western blot using anti-His and anti-MBP antibodies that detected a strong band approximately 110 kDa in size. Protein concentration was determined by Bradford assay using a BSA standard.

### Dynamic light scattering (DLS)

Aliquots of purified MBP-FUS-CHOP WT, MBP-FUS-CHOP 12A, and MBP-FUS-CHOP 12E were thawed on ice and diluted to 0.34 mg/mL. 176 µL of protein was aliquoted into two separate 0.5 mL low-bind Eppendorf tubes per protein. 20 µL of TEV protease buffer was added to all tubes. Either 4 µL of TEV protease (New England Biolabs) or 4 µL of PBS was added to each protein sample. Samples were incubated at room temperature overnight and transferred into a black wall, clear bottom 96-well plate. The plate was analyzed using a Wyatt DynaPro Plate Reader at 25°C.

### KEY RESOURCES TABLE

| REAGENT or RESOURCE | SOURCE | IDENTIFIER |
| --- | --- | --- |
| Antibodies |  |  |
| CHOP/DDIT3 | Cell Signaling Technology | #2895 |
| CHD4 | Cell Signaling Technology | #11912 |
| BRG1 | Cell Signaling Technology | #49360 |
| SNF2H | EMD Millipore | ABE1026 |
| BAF170 | Cell Signaling Technology | #12760 |
| ACF1 | Abcam | ab187670 |
| HDAC1 | Cell Signaling Technology | #5356 |
| TBP | Cell Signaling Technology | #44059 |
| H3K27me3 | Cell Signaling Technology | #9733 |
| IgG - rabbit | Cell Signaling Technology | #2729 |
| H3K27ac | Sigma | MABE647 |
| H3K27ac | Active Motif | 39133 |
| Chemicals, Peptides, and Recombinant Proteins |  |  |
| MBP-FUS-CHOP WT | This paper |  |
| MBP-FUS-CHOP 12A | This paper |  |
| MBP-FUS-CHOP 12E | This paper |  |
| Critical Commercial Assays |  |  |
| Duolink Proximity Ligation Assay | Sigma-Aldrich | DUO92101-1KT |
| Deposited Data |  |  |
| ChIP-seq and RNA-seq raw data | NCBI GEO | GSE145678 |
| Experimental Models: Cell Lines |  |  |
| NIH-3T3 | ATCC | CRL-1658 |
| SW-872 | ATCC | HTB-92 |
| MLS 402 | Gift from Pierre Aman |  |
| MLS 1765 | Gift from Pierre Aman |  |
| DL-221 | Gift from Keila Torres and Alexander Lazar |  |
| Recombinant DNA |  |  |
| N174(Puro)-FUS-CHOP WT | this paper |  |
| N174(Puro)-FUS-CHOP 12A | this paper |  |
| N174(Puro)-FUS-CHOP 12E | this paper |  |
| N174-MCS (Puro) backbone | Addgene | Addgene plasmid # 81068 ;<br>RRID:Addgene_81068 |
| Tet-pLKO-Puro | Addgene | Addgene plasmid # 21915 ;<br>RRID:Addgene_21915 |

## Supplemental Information

**Fig. S1.**
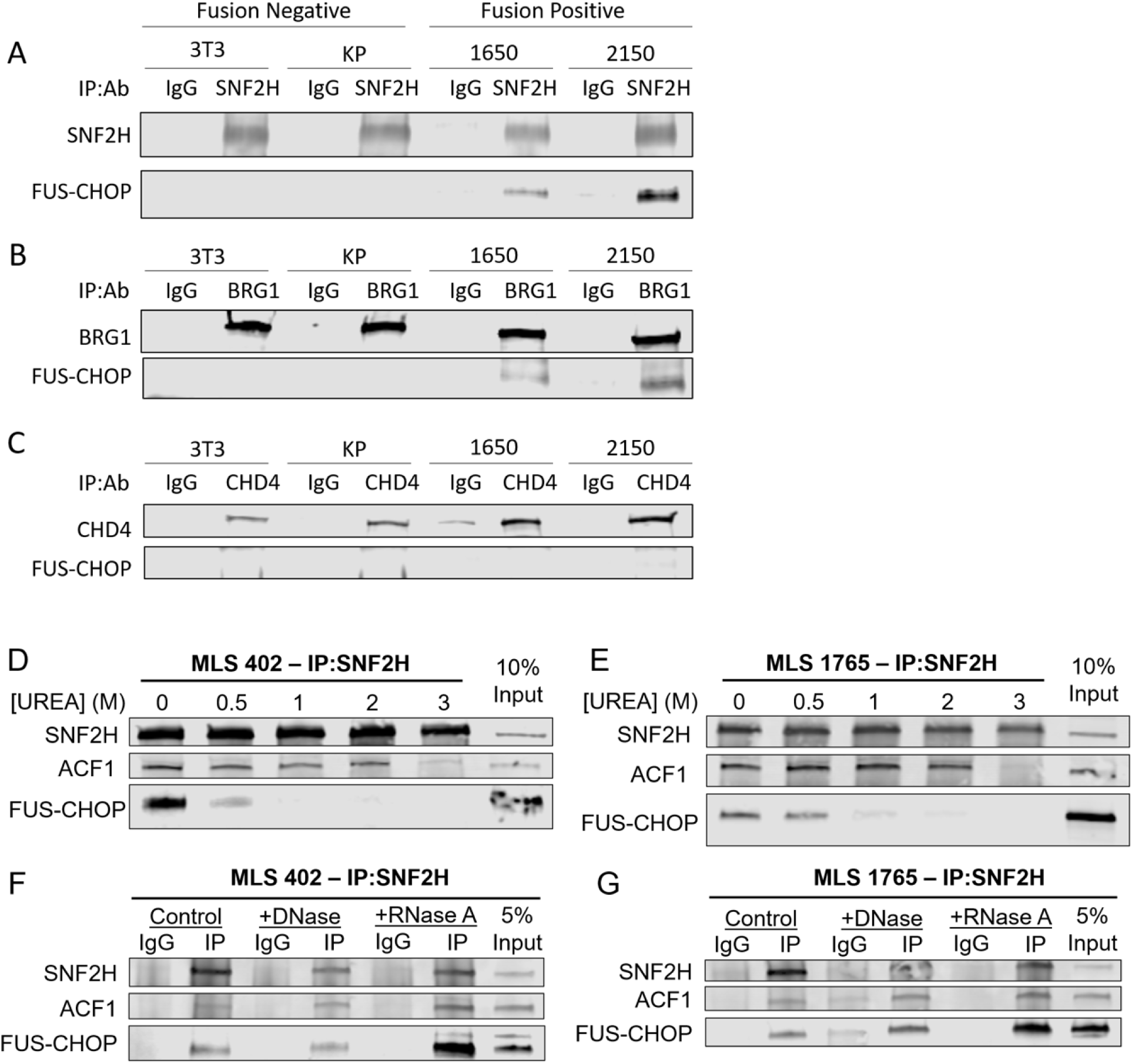
FUS-CHOP interacts with the ISWI and BAF complexes in mouse FUS-CHOP-driven sarcoma cell lines. (A) Snf2h, (B) Brg1, and (C) Chd4 co-IP for FUS-CHOP. 3T3 and KP cell lines are fusion negative. 1650 and 2150 are fusion-positive mouse sarcoma cell lines. (D) Urea denaturation assay in MLS402 and (E) MLS1765 cell lines performed with SNF2H co-IP for FUS-CHOP. ACF1 was used as a positive control for co-IP with SNF2H. (F) RNase/DNase assay in MLS402 and (G) MLS1765 cell lines performed with SNF2H co-IP for FUS-CHOP. ACF1 was used as a positive control for co-IP. Lysates were treated with either DNase or RNase A to deplete DNA and RNA in co-IP lysates, respectively.

**Fig. S2.**
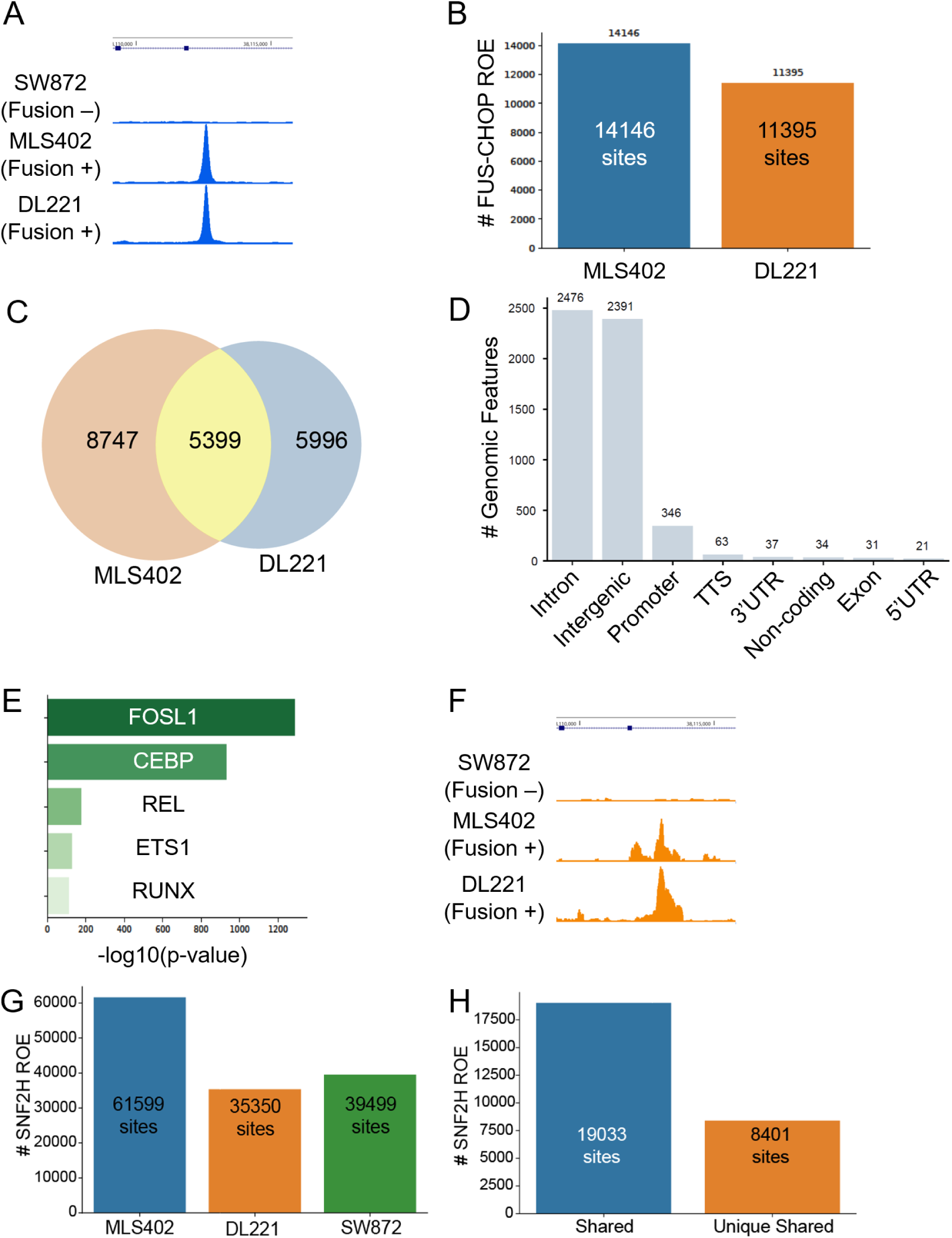
Genome-wide mapping of FUS-CHOP and SNF2H binding reveals widespread binding at enhancers and specific DNA-binding motifs in human MLPS cell lines. (A) Example track demonstrating FUS-CHOP transcription factor binding in human MLPS cell lines. (B) Total unique FUS-CHOP peaks called in MLS402 and DL221 cell lines. (C) Venn diagram showing the overlap of FUS-CHOP peaks in human MLPS cell lines. (D) Distribution of genomic features at MLS402-DL221 shared FUS-CHOP binding sites. (E) Top five DNA-binding motifs from motif analysis of unique FUS-CHOP peaks in human MLPS cell lines. (F) Example track demonstrating SNF2H binding in human MLPS cell lines. (G) Total unique SNF2H peaks called in MLS402, DL221, and SW872 cell lines. (H) Total shared SNF2H binding sites in MLS402, DL221, and SW872 cell lines, and total unique shared SNF2H binding sites in human MLPS cell lines only.

**Fig. S3.**
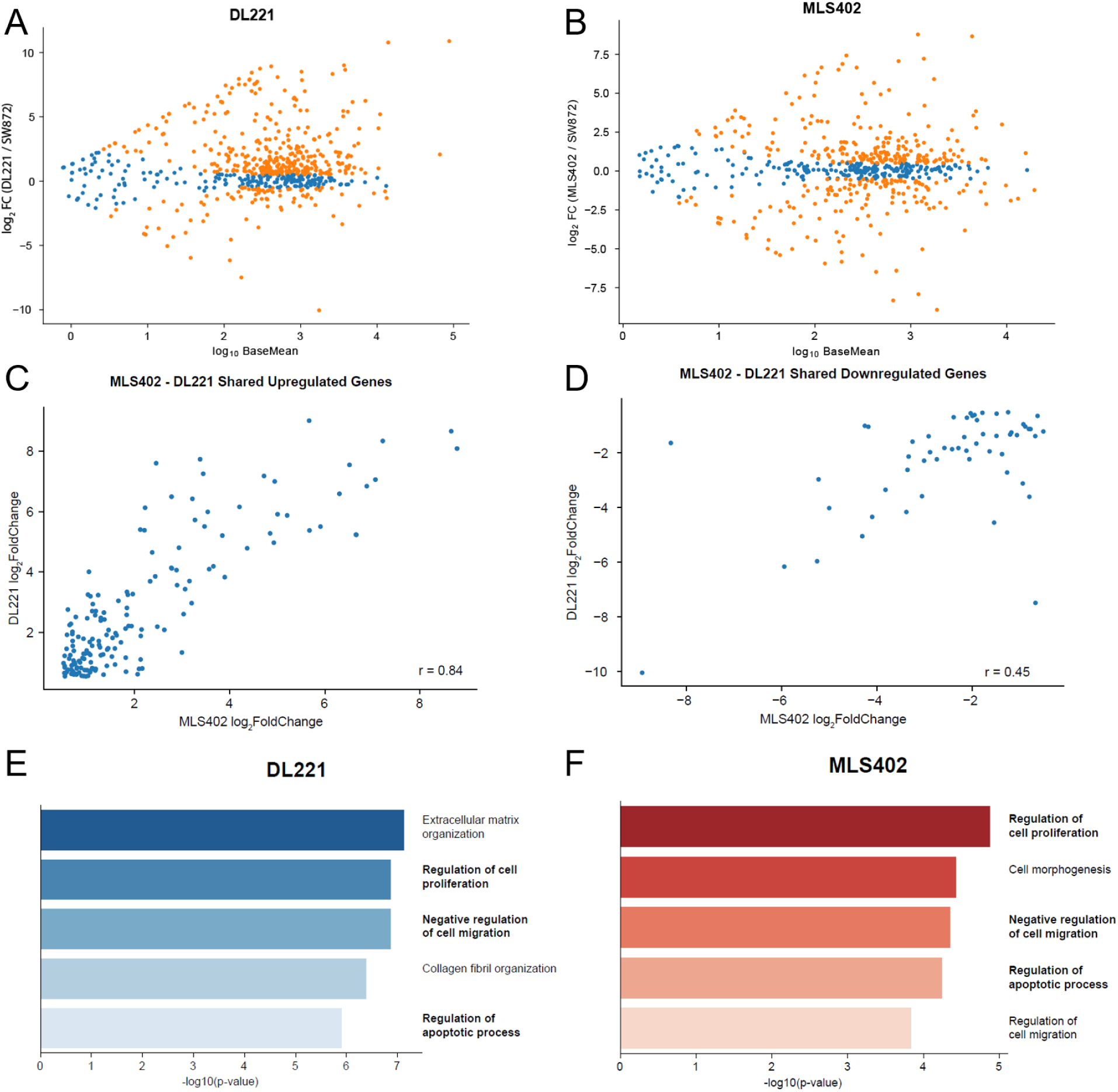
Transcriptome analysis of gene expression changes in human MLPS cells. (A) Differentially expressed genes in DL221 and (B) MLS402 MLPS cell lines compared to the SW872 liposarcoma cell line. Orange dots represent significant differential expression. Blue dots represent insignificant differential expression. (C) Scatterplot of upregulated genes in MLS402 versus upregulated genes in DL221. r = 0.84. (D) Scatterplot of downregulated genes in MLPS402 versus downregulated genes in DL221. r = 0.45. (E) Enrichr gene ontology (GO) biological process results for upregulated genes in DL221 and (E) in MLS402 human MLPS cell lines.

**Fig. S4.**
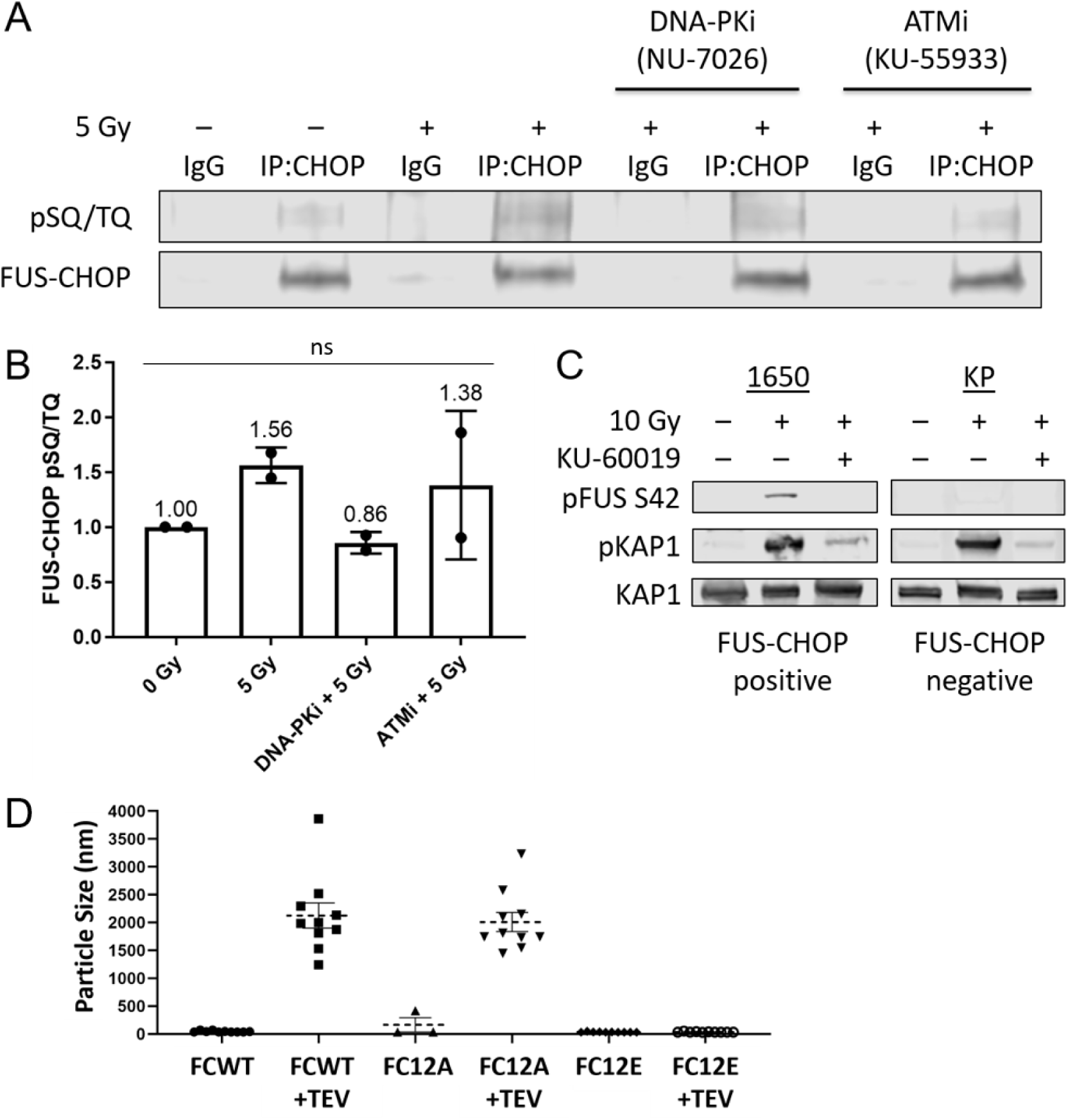
FUS-CHOP is phosphorylated after irradiation by DNA-PK and ATM. (A) IP of FUS-CHOP from MLS402 cell lines treated with no radiation, 10 Gy X-ray radiation, NU-7026 with radiation, and KU-55933 with irradiation. (B) Quantification of FUS-CHOP phosphorylation detected on immunoblot using a phospho-SQ/TQ antibody. ns = not significant, One-way ANOVA with multiple comparisons. (C) Immunoblot of phosphorylation at Ser42 in FUS-CHOP after irradiation and treatment with an ATM inhibitor, KU-60019. (D) FUS-CHOP (FCWT), FUS-CHOP 12A (FC12A), and FUS-CHOP 12E (FC12E) particle size before and after the addition of TEV protease to release the MBP solubility tag.

